# Unveiling the intertwined roles of the spatial-temporal environment and behavioural modes in animal movement

**DOI:** 10.1101/2022.12.19.521042

**Authors:** Hans Linssen, Henrik J. de Knegt, Jasper A.J. Eikelboom

## Abstract

1. Animal movement arises from complex interactions between animals and their heterogeneous environment, making it challenging to capture the movement process in a single analysis. This has lead to a bias towards lower-dimensional representations of the process in such analyses. In order to better understand animal movement, its multiple components should be included and addressed simultaneously.
2. We present an analytic framework that integrates the *behavioural*, *spatial*, and *temporal* components of the movement process and their interactions, and allows for assessing the relative importance of these components. We propose a *daily cyclic covariate* to represent temporally cyclic movement patterns, for example diel variation in activity, and combine the three components in multi-modal Hidden Markov Models. We compare the statistical fits of models that include or exclude any of the behavioural, spatial and temporal components, and perform variance partitioning on the model that included all components to assess their relative importance to the movement process, both in isolation and in interaction.
3. We apply our framework to a case study on the movements of zebra, wildebeest and eland antelope in a South African reserve. Behavioural modes impacted movement the most, followed by diel rhythms and then the spatial environment (i.e. tree cover and terrain slope). Interactions between the components often explained more of the movement variation than the marginal effect of the spatial environment did on its own. Omitting components from the analysis led either to failure to detect relationships between input and response variables, resulting in overgeneralisations when drawing conclusions about the movement process, or to erroneously detecting spurious relationships, resulting in factually incorrect conclusions.
4. Our analytic framework can be used to study animal movement by integrating the different components of the movement process, thereby preventing incomplete or overly generic ecological interpretations. We demonstrate that understanding the drivers of animal movement, and ultimately the ecological phenomena that emerge from it, critically depends on considering the various components of the movement process, and especially the interactions between them.

Data/code for peer review statement: the data and code needed to reproduce the study and results are uploaded as a file with the initial submission.

## 1 Introduction

Movement is ubiquitous to animal life and central to emergent ecological phenomena such as territoriality and home range formation (Kie et al., 2002; Moorcroft et al., 1999), predator-prey dynamics (Abrams, 2007; Flaxman & Lou, 2009), distribution patterns (Bailey et al., 1996) and population dynamics (Morales et al., 2010; Revilla & Wiegand, 2008). To understand such phenomena it is thus key to study what drives the movement of animals through their environment (Morales et al., 2010; Schick et al., 2008). However, this understanding is hampered by the complex relationships between animal movement and a multitude of behavioural and spatial-temporal environmental drivers, since the same state of a single driver can give rise to different movement patterns depending on the state of other drivers. For example, animals might move slowly through thick vegetation when foraging, or fast if they are translocating from one preferred high-visibility habitat to another. Temporal variation in the response of animals to their environment in diel, lunar or seasonal patterns (e.g. Riotte-Lambert et al., 2013), complicates the analysis of animal movement drivers even further. In order to structure the study of animal movement and advance the interpretation of movement patterns, a movement ecology framework has been put forward that divides the movement process into multiple components: an internal motive regarding *why* to move and the temporally and spatially varying environment influencing *when* and *where* to move respectively (Nathan et al., 2008).

First, ‘*why move*’ relates to the internal motives of movement (e.g. reproduction, resource acquisition and threat avoidance), which are often manifested in different behaviours (e.g. mating, foraging and fleeing) that produce distinct movement modes (i.e., movement trajectory geometries that can be quantified by metrics like speed and turning angle distributions) (McClintock et al., 2012; Morales et al., 2004; Nathan et al., 2008). Distinguishing different movement modes in animal trajectories has yielded insights in different foraging strategies within individuals (McClintock et al., 2017), predator-prey interactions (Forester et al., 2007) and effects of anthropogenic disturbance on behaviour (van Beest et al., 2019). Internal motives and their emergent movement modes should thus be acknowledged to further the interpretation of animal movement, specifically regarding its iteraction with the environment (Jonsen et al., 2006; McClintock et al., 2012, 2017; Morales et al., 2004).

Second, *‘when to move’* deals with temporal variation in animal movement, which often exhibits cyclic patterns. For example, seasonal migratory movements (e.g. Dingle, 2014; Dingle & Drake, 2007) have long been recognized, as well as the influence of lunar rhythms (e.g. Polansky et al., 2010; Riotte-Lambert & Matthiopoulos, 2020). Moreover, most animals show strong diel activity patterns (Ensing et al., 2014; Green & Bear, 1990; Owen-Smith & Goodall, 2014; Vazquez et al., 2019), which can arise directly due to variation in light and thermal conditions (Dussault et al., 2004) or indirectly due to predation risk and human activity (Ensing et al., 2014; Fischhoff et al., 2007; Owen-Smith & Traill, 2017; Skarin et al., 2010). Temporal rhythms are thus a key characteristic of animal movement (Riotte-Lambert et al., 2013).

Third, *‘where to move’* relates the movement of animals to the characteristics of their spatial surroundings. For example, topography influences movement through energetic costs associated with moving through sloping or rugged terrain (Dailey & Hobbs, 1989; Parker et al., 1984; Wall et al., 2006; White & Yousef, 1978). Daily cycles can interact strongly with the spatial surrounding, for example in animals that select areas with high vegetation cover during daytime and more open areas during the night to decrease predation risk (Ager et al., 2003; Leblond et al., 2010). These examples illustrate that relationships between spatial environmental characteristics and movement patterns are conditional on the context of internal motive and time, as this context determines the costs and benefits that the spatial environmental characteristics represent (e.g. Lubitz et al., 2022).

Internal motives, temporal variation and spatial environmental heterogeneity are thus three intricately linked components of the animal movement process. However, animal movement research has often been hampered by the focus on univariate analyses (Fig. 1; Eikelboom et al., 2020), while ideally the relationships between animal movement and each of these three components are considered within each other’s context (Nathan et al., 2008). Although the movement ecology framework has proven helpful in elucidating drivers of animal movement (Joo et al., 2022), challenges remain regarding how to 1) include these three components in a single coherent model framework, 2) quantify the relative contributions of the components in shaping animal movement, and 3) assess the influence of interactions between the components on the quantitative and qualitative inferences made about the movement process.

**Figure 1.**
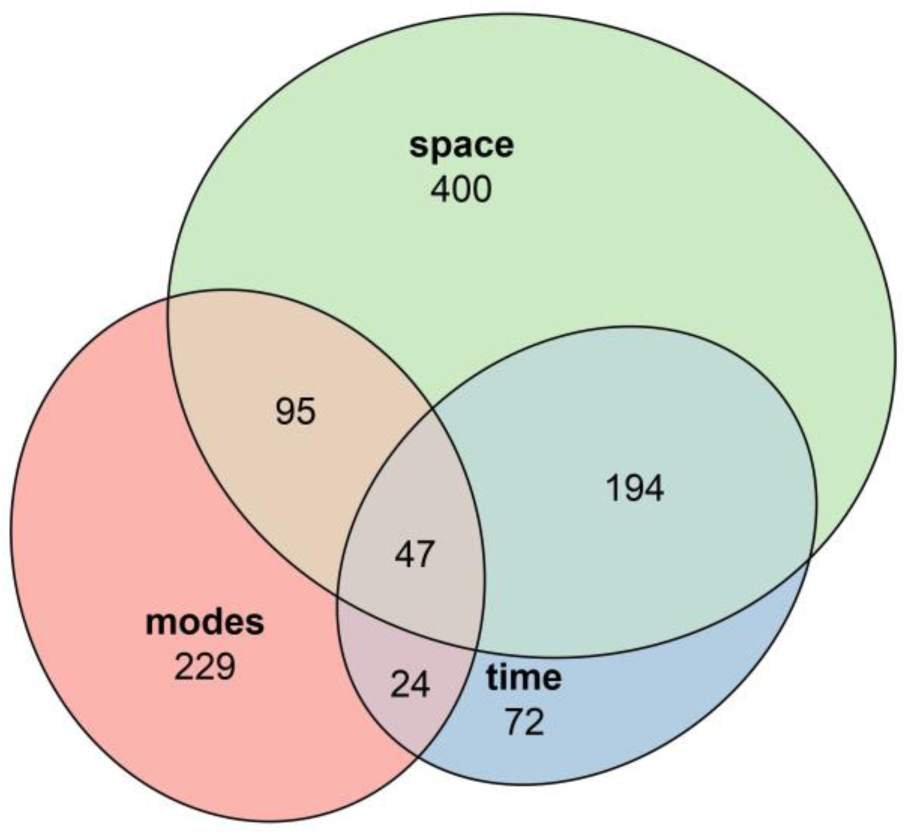
Results of a systematic literature search (Scopus, 16-03-2023; see Supporting Information), demonstrating the numbers of studies that focused on the different components of the animal movement process (modes, time and space) separately or combined.

Here, we demonstrate an integrative analytic framework to study the role of movement modes, time (temporal rhythms) and space (environmental heterogeneity) in shaping animal movement. We apply this framework to a case study of large savanna ungulates. We quantify the relative contribution of these three components to the movement process in isolation as well as in interaction, and we assess how ecological interpretations of movement change when omitting these interactions from analyses. We use fine-scale (10-minute resolution) GPS tracking data of eland antelope (*Taurotragus oryx*), blue wildebeest (*Connochaetes taurinus*) and plains zebra (*Equus quagga*) in a South-African game reserve, and fit Hidden Markov Models (HMMs) with input from tree cover, terrain slope and a newly developed *daily cyclic covariate*. Since HMMs can describe the movement process as a multi-modal random walk with input from spatial-temporal environmental covariates (McClintock et al., 2020), they provide an ideal method to quantify the effects of modes (as a proxy for internal motives) and spatial and temporal environmental heterogeneity on movement. Besides setting forth an integrative analytic framework to study animal movement in light of all three movement process components simultaneously, we use our case study of savanna ungulates to demonstrate that inferences about the behavioural ecology of animals are made best when explicitly modelling the movement process with these three components in interaction.

## 2 Materials and methods

### 2.1 Animals and study area

From June to August 2017, 34 eland antelope, 34 blue wildebeest and 35 plains zebra were captured in Welgevonden Game Reserve, Limpopo, South Africa (24°13’S, 27°54’E), and equipped with neck collars containing GPS and accelerometer sensors. For more details about the collaring process and sensors, see de Knegt et al. (2021). The procedures were approved by Welgevonden Game Reserve as a management action and carried out in accordance with relevant guidelines and regulations under the National Environmental Management Protected Areas Act (NEM: PAA; Act No. 57 of 2003; see Supporting Information). The animals were released in a fenced study area of 1200 hectares at the northern edge of the reserve (Fig. 2). As a part of the Waterberg mountain massif and Bushveld ecoregion (Mucina et al., 2006), the study area is characterised by nutrient poor sandy soils in the lower northern part and rocky, flat hilltop plateaus dissected by ravines extending towards the south. Woody vegetation varies from mixed to open dry deciduous woodland, with the rocky plateaus being the most open.

**Figure 2.**
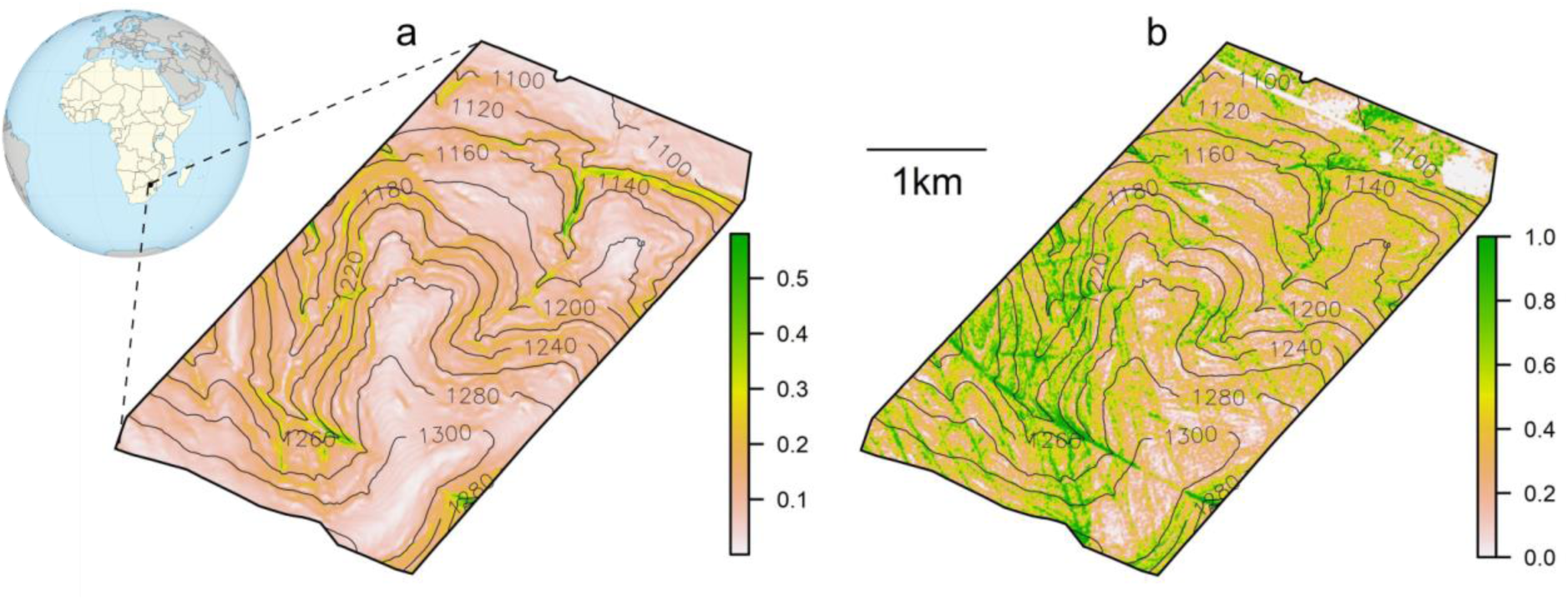
Overview of the study area with a) terrain slope (radians) and b) fractional tree cover. Numbers at the isoclines denote elevation from sea level (m).

### 2.2 Animal movement data

The GPS sensors collected positions with an interval ranging between 15 and 2 minutes, depending on the amount of body activity as determined by the accelerometer (de Knegt et al., 2021). The position data were corrected, smoothed and modelled to regular 10 min resolution trajectories as described in de Knegt et al. (2021). Due to sensor failures, we selected from each species only the ten individuals with the most recorded data in the period from September to December 2017 (the first four months of the study period, during which the sensors collected most data). Our subset contained a total of 357,815 animal positions across 30 individuals of the three species, comprising 481 bouts (i.e. an uninterrupted and regular sequence of positions of a single individual) with a median bout length of 496 positions (3.4 days).

### 2.3 Environmental data

A digital elevation model of the study area with resolution 2×2 m was available, from which terrain slope (hereafter referred to as slope) was calculated (Fig. 2a). We used aerial images of the study area from August 2013 to create an orthomosaic with a 0.25×0.25 m resolution, from which pixels were classified into trees, grass, bare ground and other/built up area using the Semi-Automatic Classification Plugin for QGIS (Congedo, 2021). After having applied an isotropic Gaussian kernel smoother with a 10 m standard deviation to the classified pixels, we derived fractional tree cover (hereafter referred to as tree cover) and fractional grass cover maps of the study area at a 2×2 m resolution (Fig. 2b). Tree cover was strongly negatively correlated with grass cover (*r* = 0.95), so we used only tree cover in our analyses and considered it a measure for the inverse of grass cover as well. We extracted tree cover and slope values for each animal position.

### 2.4 Movement model structure

We constructed a Hidden Markov Model (HMM) for each species separately to examine how their movement was shaped by movement modes, diel patterns and environmental heterogeneity. We described the movement process as a random walk consisting of steps between subsequent animal positions with lengths *s* and turning angles *φ*. Since intervals between steps were constant at ten minutes, we refer to *s* as speed throughout. *s* and *φ* were fitted to *gamma* (*Γ*) and *von Mises* (*vM*) distributions, respectively, which were unique to each discrete movement mode *z*. Thus, *z* was defined as a unique combination of three movement parameters: the mean *μ* and standard deviation *σ* of the speed (converted from the shape and scale parameter of the *Γ* distribution for easier interpretation) and the *vM* concentration parameter *κ* of the turning angle (the *vM* mean was fixed at 0 since there was no reason to assume a movement bias to either the right or left direction). *μ*, *σ* and *κ* were in turn modelled as functions of several covariates (see section *Movement model covariates*) using a log-link. Mode switching between the steps was governed by a switching matrix *M*, where the off-diagonal elements are logit-linear functions of covariates. We considered a three-mode, two-mode and a ‘single-mode’ HMM (i.e. 3×3, 2×2 and 1×1 dimensions for *M* respectively), where the single-mode model is analogous to a movement model that does not include the mode component. Thus, the HMM had the general form:

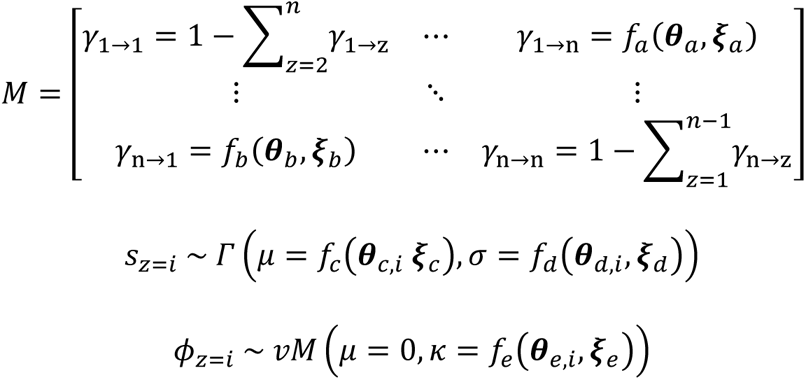

Where *γ_i→j_* is the probability of switching from mode *z = i* to *z = j* between consecutive steps, *f_n_* are functions for step switching probabilities and *Γ* and *vM* parameters, ***ϑ****_n_* are sets of to-be-fitted coefficients of function *f_n_* (and for mode *z = i* in case of ***ϑ****_n,i_*), and ***ξ****_n_* are sets of covariates (see next section).

### 2.5 Movement model covariates

We modelled the parameters *μ*, *σ* and *κ* and the mode switching probabilities *γ_i→j_* each as a function of three covariates. Two of those were the spatial environmental covariates tree cover and slope. The third covariate was a constructed *daily cyclic covariate* (DCC) that describes the diel patterns in *μ*, *σ* and *κ* (driven by e.g. ambient temperature, solar position, perceived predation risk and internal diel rhythm). To determine those diel patterns, we binned the animal positions per time of day (144 bins of 10 minutes) and fitted *Γ* distributions (*μ* and *σ*) to speeds and *vM* distributions (*κ*) to turning angles within a centred moving window of three bins wide. This yielded empirical diel patterns in parameters *μ*, *σ* and *κ* (Fig. S1). We constructed DCC by first rescaling and averaging the empirical diel patterns in *μ*, *σ* and *κ* to one aggregate diel pattern per species (Fig. 3), given that these separate empirical diel patterns were strongly correlated within species (Fig. S2). We then fitted to that aggregate diel pattern a mixture of a cosine and four wrapped normal distributions with 24 h periods to compute the final DCC values per species, with the wrapped normal distributions representing two peaks and two lows per day that were fixed at the peaks and lows of the aggregate diel patterns (Fig. 3). Thus, DCC had the following form:

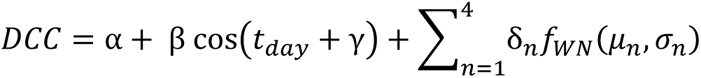

**Figure 3.**
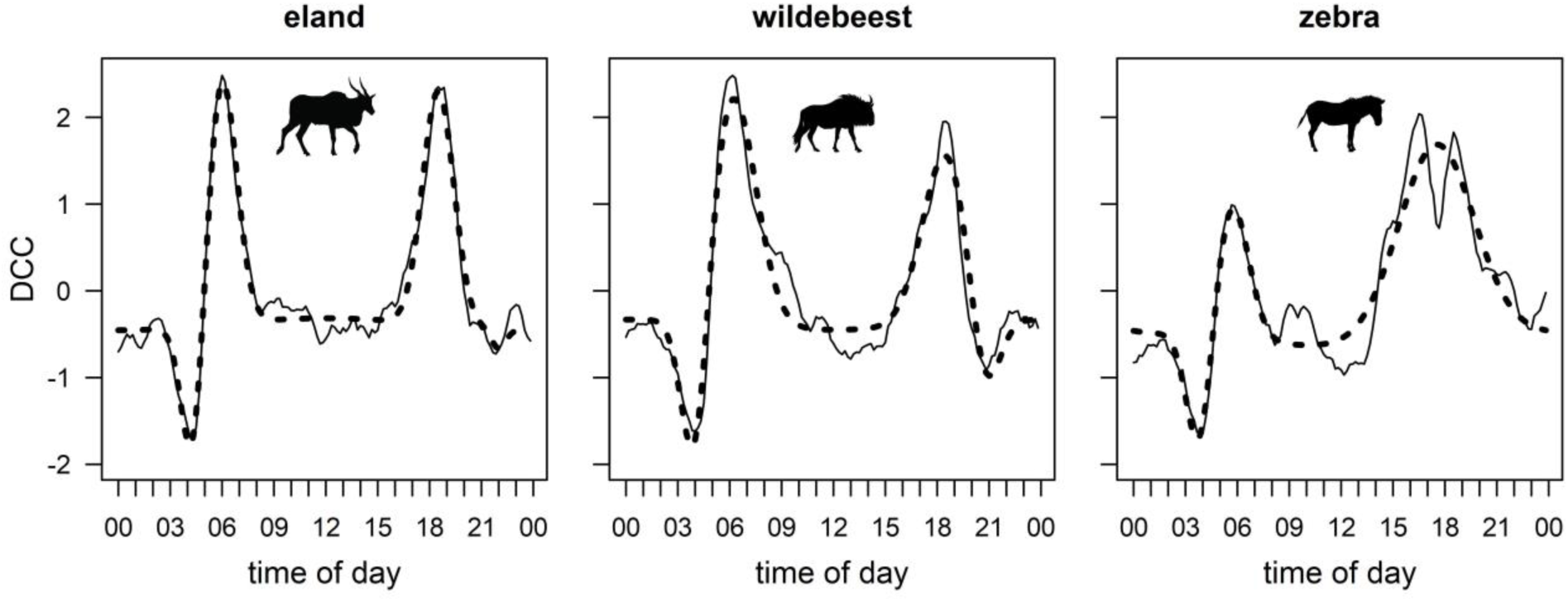
Aggregate diel patterns averaged from *μ*, *σ* and *κ* for eland, wildebeest and zebra (solid line). We fitted a daily cyclic covariate (DCC, dashed line) to those aggregate diel patterns, which summarizes the diel activity patterns per species as expressed in the movement properties *μ*, *σ* and *κ*. DCC is a mixture distribution of a cosine and four wrapped normal distributions with 24 h periods.

Where *t_day_* is the time of day (range 0 – 2π), *f_WN_* is the probability density function of the wrapped normal distribution with a 24 h period (describing 1 out of 4 peaks or lows per day in the aggregate diel pattern), and *δ_n_* are scalars for those distributions. As a result, each animal position had a corresponding DCC value, which depended on the time of day and the species (Fig. 3). We then standardized tree cover, slope and DCC to zero-mean and unit variance, and included the three variables and their two-way interactions as covariates of *μ*, *σ* and *κ* in the HMMs. We included tree cover, slope and DCC without interactions as covariates of the mode switching probabilities *γ_i→j_*.

### 2.6 Model fitting workflow

We fitted a series of HMMs per species separately. The HMM series consisted of models containing any combination of the three components of the movement process (modes, time and space), ranging from a null model that contained none, to a full model that contained all three in interaction (Table 1). The models were fit by numerical maximisation of the log-likelihood function, using the *fitHMM* routine from the R package *momentuHMM* (McClintock & Michelot, 2018). We fitted the models in order of increasing complexity, every step adding covariates and/or interactions between covariates to the distribution parameters and switching probabilities. The full model included all three movement components modes, time and space, namely: DCC, tree cover and slope as covariates to the mode switching probabilities *γ_i→j_*, and these three variables and their two-way interactions as covariates to *μ*, *σ* and *κ*. The fitted parameter values from each fitting step were used as starting values for the next; starting values for other parameters were set to 0 (note that all covariates had been scaled to mean 0 and standard deviation 1). For each iteration, four sets of perturbations of those starting values *β* were sampled from ∼*N*(*β*, *σ* = 0.25 · *β*). The model fit with the highest likelihood out of all starting sets was the optimal fit of that iteration, and it was used to initiate the model fit of the next iteration. However, in practice all iteration sets of fitted models converged to almost identical coefficient values and model likelihoods. We used the Akaike information criterion (AIC; Akaike, 1973) to assess model parsimony throughout the workflow. The model structure and fitting workflow were identical for all three species.

**Table 1.** Overview of the fitted HMMs, each including a combination of the predictors modes, time and space. ΔAIC was calculated for each species separately relative to the best model of that species. The AIC of the full models (model 3) was 1,305,636; 1,304,360 and 1,286,972 for eland, wildebeest and zebra respectively. tc = tree cover, sl = terrain slope, DCC = daily cyclic covariate, *μ*, *σ* and *κ* are movement parameters and *γ_i→j_* = mode switching probability. Powers of 2 indicate all two-way interactions among the covariates between brackets.

| Model | Predictor set | #modes | $\mu, \sigma$ and $\kappa$ | $\gamma_{i \rightarrow j}$ | $\Delta AIC$ | $\Delta AIC$ | $\Delta AIC$ |
| --- | --- | --- | --- | --- | --- | --- | --- |
|  |  |  | covariates | covariates | eland | wildebeest | zebra |
| 0 | none | 1 | – | – | 85,474 | 64,152 | 57,309 |
| 1a | modes | 2 | – | – | 10,932 | 8,974 | 7,572 |
| 1b | time | 1 | DCC | DCC | 74,877 | 49,193 | 50,430 |
| 1c | space | 1 | (tc, sl)<sup>2</sup> | tc, sl | 84,674 | 63,346 | 55,937 |
| 2a | modes + time | 2 | DCC | DCC | 822 | 918 | 2,240 |
| 2b | modes + space | 2 | (tc, sl)<sup>2</sup> | tc, sl | 10,157 | 8,219 | 5,327 |
| 2c | time + space | 1 | (tc, sl, DCC)<sup>2</sup> | tc, sl, DCC | 74,117 | 48,110 | 48,601 |
| 3 | modes + time + space | 2 | (tc, sl, DCC)<sup>2</sup> | tc, sl, DCC | 0 | 0 | 0 |

### 2.7 Variance partitioning of movement process components

Based on the fitted HMM with all predictors included (model 3 in Table 1), we partitioned the explained variance of the modelled parameters *μ*, *σ* and *κ* for each species into contributions from the three components of the movement process: modes, time and space. To quantify what proportion of the total variance in *μ*, *σ* and *κ* was accounted for by each of the three components, we used linear regressions with the predicted values for *μ*, *σ* and *κ* as response variables (on the log-scale) based on the most probable movement mode, and the three components as predictors. The most probable movement modes along the movement path were determined using Viterbi decoding (Zucchini & Macdonald, 2009). We alternately omitted or included the mode component (i.e. a single-mode or multi-modal model with the assigned highest mode probabilities), the time component (DCC), the space component (tree cover and slope) or their interactions as predictors and summarised the regression fit using *R^2^* values. By doing so, we could partition the variance of the fitted model parameters *μ*, *σ* and *κ* into the independent model components (i.e. mode, time and space), as well as their combined effects. All analyses were performed in R 3.6.1 (R Core Team, 2023).

## 3 Results

### 3.1 Hidden Markov Models

We fitted Hidden Markov Models (HMMs) that included combinations of the three components of the movement process: modes, time and space (Table 1). When including the modes component (Models 1a, 2a, 2b and 3), we explored models with two and three movement modes. The full three-mode models achieved a higher model fit than the full two-mode models, albeit with a relatively small difference (ΔAIC = 32,093; 29,170 and 27,381 for eland, wildebeest and zebra respectively). Both the two- and three-mode models detected one faster, more directed mode and one slower, more tortuous mode (Fig. S3). We labelled these as ‘transit’ and ‘encamped’ modes, considering the labels to be reasonable abstractions while retaining spatially descriptive meaning and avoiding to project too specific behavioural annotations on the modes. The three-mode HMMs detected an additional slow and directional mode (Fig. S3). This third mode responded relatively weak to the temporal and spatial environmental covariates, while the response displayed by the other two modes was qualitatively similar for both the two- and three-mode HMMs (Figs. S4-S6). Therefore, we do not further consider the three-mode HMMs, and hereafter refer to the two-mode HMMs when discussing models that include movement modes.

The AIC values of all HMMs are presented in Table 1. The inclusion of movement modes led to much higher model fits, with the models including modes but not time and space (Model 1a) outperforming the models including time and space but not modes (Model 2c; ΔAIC = 63,185; 39,136 and 41,029 for eland, wildebeest and zebra respectively). Furthermore, the models including time but not space were superior to those including space but not time (Model 1c versus 1b; ΔAIC = 9,797; 14,153 and 5,507 for eland, wildebeest and zebra respectively; Model 2b versus 2a; ΔAIC = 9,335; 7,301 and 3,087 for eland, wildebeest and zebra respectively), indicating that DCC is a dominant covariate accounting for substantial temporal variation, which is not accounted for by tree cover and slope. For each species the full model, which includes modes, time and space (Model 3), was the most parsimonious model according to AIC value.

### 3.2 Variance partitioning of movement process components

Partitioning the variance in the modelled parameters *μ*, *σ* and *κ* for each species between the modes, time and space components showed that modes explained most of the variation (see Fig. S7). The contributions of the model components to the *within-mode* variation in model parameters for the three species are shown in Figure 4. For most combinations of species, parameter and modes, the temporal component explained most of the variance. However, the interactive effects of spatial-temporal predictors contributed notably to the variance in *κ*, especially for zebra (Fig. 4). By themselves, the spatial predictors did not explain much variation in movement parameters except for *κ* in zebra, and *σ* in eland (encamped mode) and zebra (transit mode; Fig. 4).

**Figure 4.**
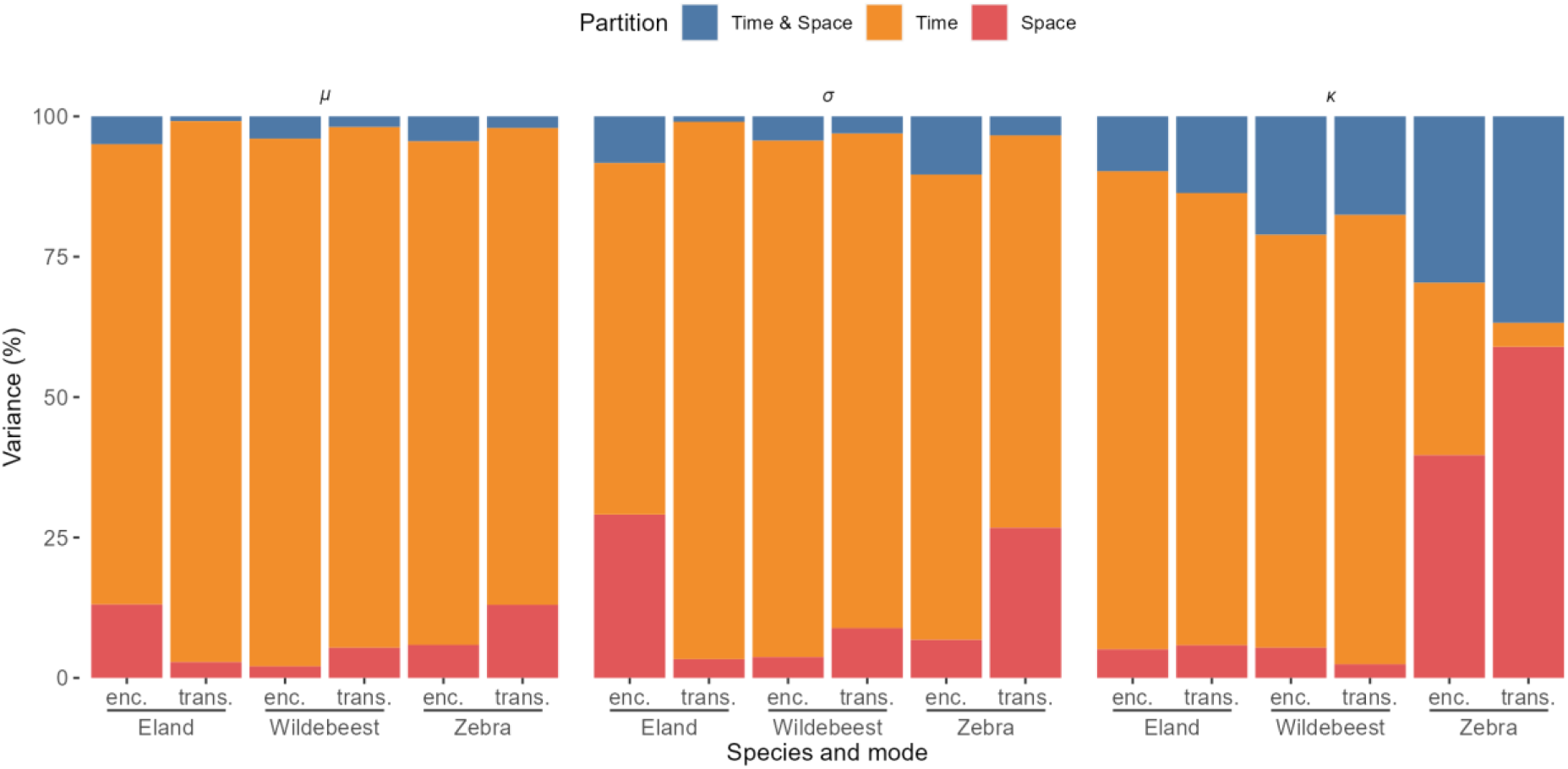
Variance partitioning, separately per movement parameter, species and movement mode (enc.= encamped, trans = transit). For variance partitioning that includes modes as a partition, see Fig. S7.

**Figure 5a.**
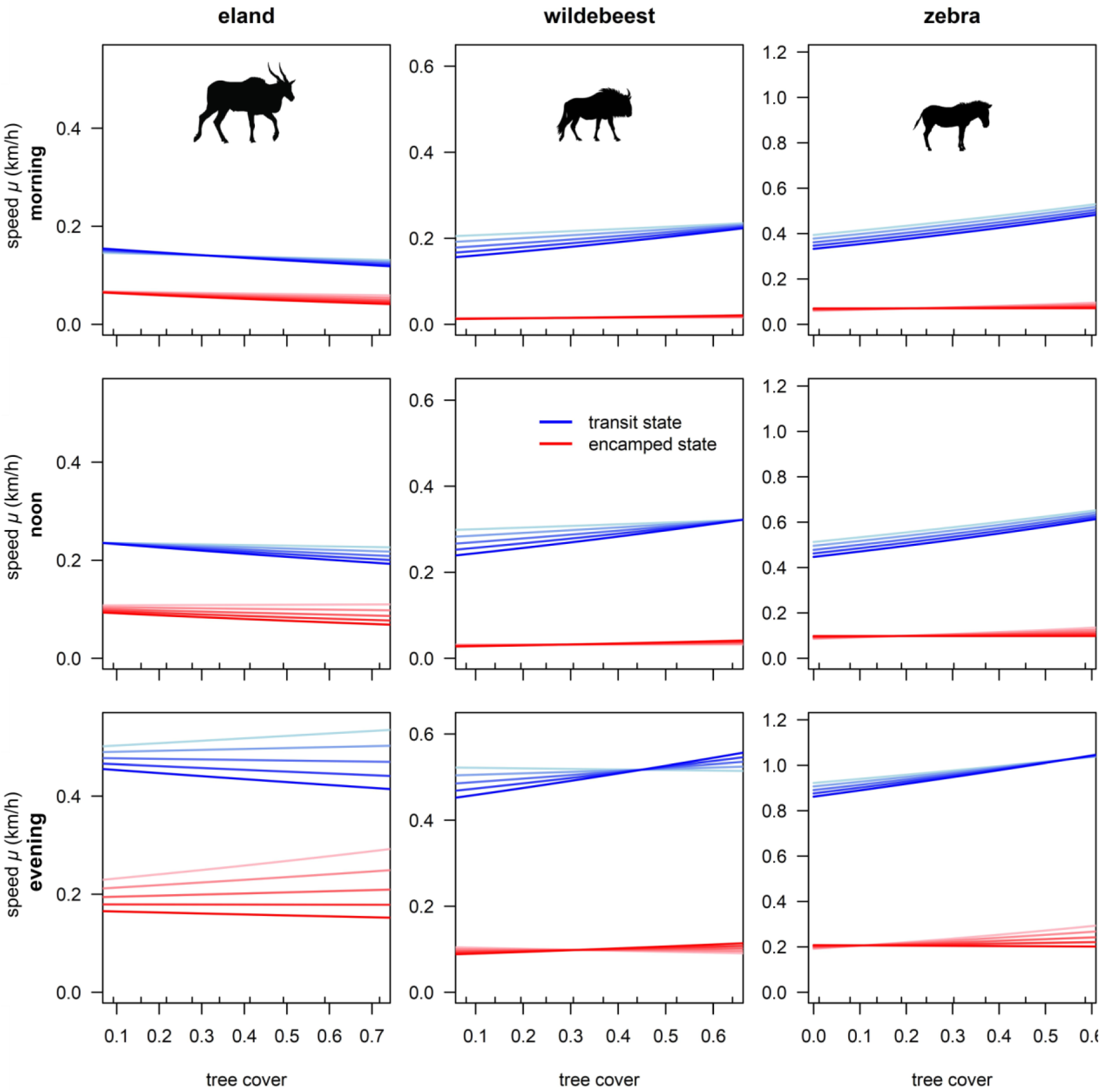
Model predictions of movement speed *μ* as a result of tree cover and slope in the morning (04:00, top row), at noon (12:00, middle row) and in the evening (18:00, bottom row) for eland, wildebeest and zebra. The times were chosen to capture a dip, baseline and peak in DCC (see Fig. 3). Blue and red indicate the transit and encamped mode respectively. Colour mapping within blue and red indicates a progression of slope from 0 (light) till 0.15 (the 90th percentile of slope values in the complete dataset; dark). Small ticks at the inside of tree cover axes mark the 10th, 20th … 90th percentile values of tree cover in the respective data sets. Note that the y-axis scales differ between species, but not between the times of day. Confidence intervals have been omitted for visual clarity, but these were generally small.

**Figure 5b.**
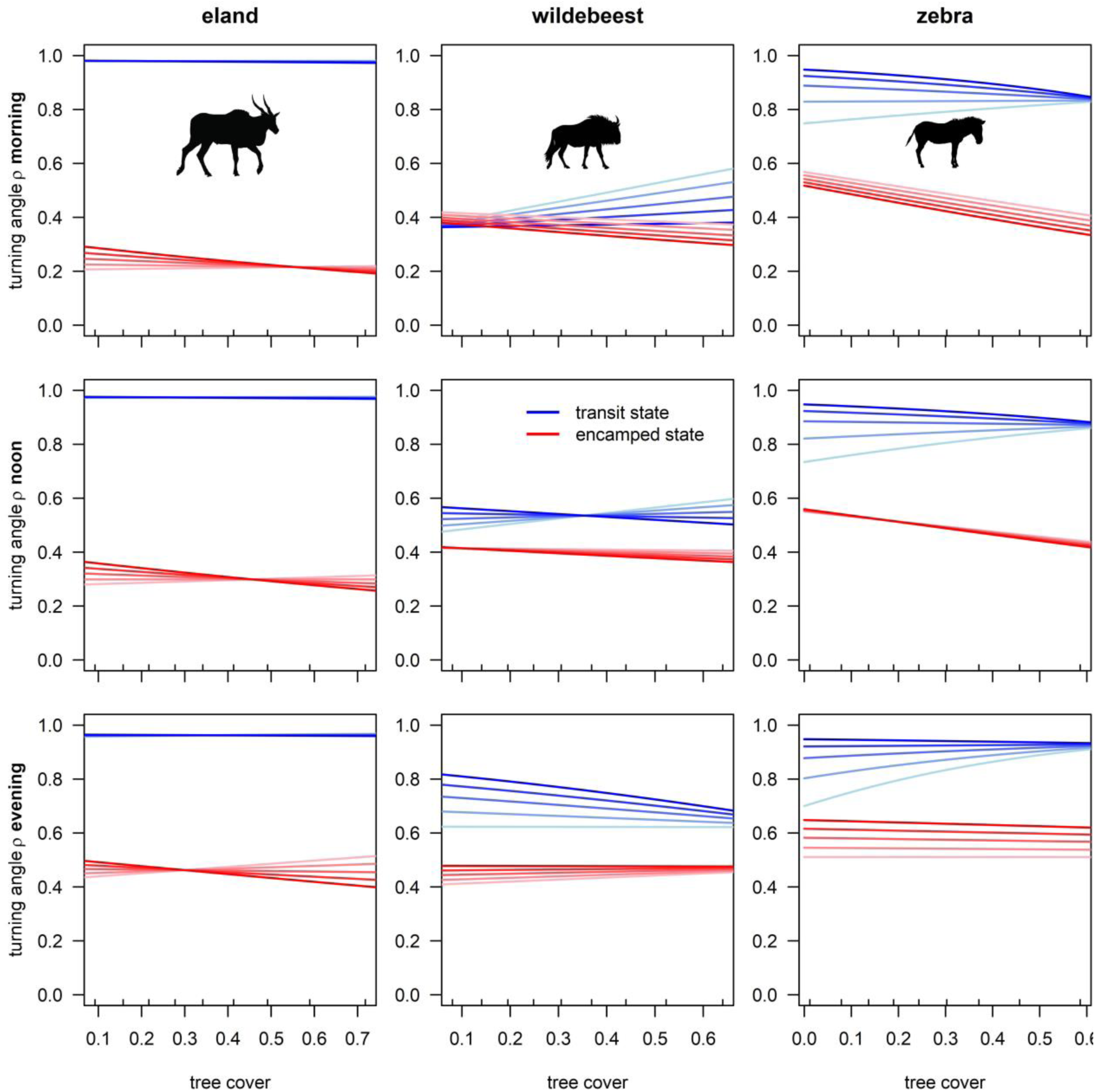
Idem to Fig. 5a, instead with model predictions of turning angle concentration on the y-axis. Concentration parameter *κ* (range 0-∞) has been converted to *ρ* (range 0-1) in this figure for visualisation purposes.

### 3.3 Different model descriptions of the movement process

Incrementally expanding the predictor set along the lines of Table 1 (Models 0 to 3) yielded various interpretations of the movement process (Figs. S8-S14). Model 0, which included neither space, time nor mode as predictors, showed that wildebeest moved on average slowest and most tortuous, with eland and zebra moving equally directed and zebra fastest (Fig. S8). When we included movement modes (Model 1a), zebra showed the same speed as eland in the encamped mode and wildebeest moved faster than eland in the transit mode, instead of being slower overall (Fig. S9). Model 1b, which included only time as a predictor, showed large fluctuations in speed and tortuosity throughout the day, with both movement parameters peaking around sunrise and sunset and showing lows before sunrise and after sunset (Fig. S10). Model 1c, including only spatial predictors, showed that speed decreased with tree cover for eland, but increased for the other species (Fig. S11). Furthermore, speed decreased with slope irrespective of tree cover. Tortuosity decreased with tree cover but increased very slighty with slope. When we included both mode and time (Model 2a), we found the same temporal patterns in speed and tortuosity as the model including only time, but also indications that daily peaks in speed were much higher for animals in the transit mode than those in the encamped mode (Fig. S12). Model 2b, including both modes and space as predictors, showcased that often only in one of the two modes did animals respond substantially to spatial heterogeneity (Fig. S13). For example, only in the transit modes did animals strongly decrease (eland) or increase (wildebeest and zebra) their speed. Zebra did not respond to slope, whereas the model including only spatial predictors suggested it slowed down on sloping terrain. Including space and time as predictors (Model 2c) reveiled that tree cover, slope and time of day interacted in their effect on speed and tortuosity, with their effects being most pronounced in the evening (Fig. S14). Model 3, including space, time and modes all as predictors, yielded the most complete description of movement, with aspects that were not apparent from all previous models (Figs. 5a,b). The movement properties displayed a strong diel pattern, with the effects of tree cover and slope on speed being largest in the evening when animals generally moved faster. Wildebeest and zebra in their transit mode moved faster with increasing tree cover whereas eland moved slower, but only so on steep terrain. In the encamped mode, tree cover had almost no effect on speed, but a strong effect on tortuosity. The animals also generally moved more tortuously with higher tree cover, even more so on sloping terrain. The model indicated that the spatial or temporal environment did not impact movement in all modes, and that the interaction effect between slope and tree cover on movement was also mode-specific.

## 4 Discussion

We developed an integrative analytic framework to quantify and understand the roles of movement modes (‘*why move*’), temporal rhythms (‘*when to move*’) and spatial environmental heterogeneity (‘*where to move*’) in shaping animal movement. In this framework, multi-modal (*why*) Hidden Markov Models (HMMs) are fitted on animal movement data, with spatial environmental covariates (*where*) and our newly developed *daily cyclic covariate* (*when*, which aggregates the empirical diel patterns in the animals’ movement parameters) influencing both the movement parameters and the mode-switching probabilities. Finally, by using a variance partitioning approach on the modelled parameters, we quantified the relative importance of the *why*-, *when*- and *where*-components and their interactions in shaping animal movement.

We demonstrated the usefulness of our analytic framework by applying it on a case study of African ungulates. The variance partitioning showed that movement modes impacted movement the most, followed by diel rhythms and the spatial environment (i.e., tree cover and terrain slope) being the least important. Furthermore, the interactions between these components were important contributors as well, often explaining more of the movement variation than the marginal effect of the spatial environment did. This contrasts recent movement ecology research, where most studies seem to focus on the spatial environment without considering movement modes and temporal rhythms (Fig. 1). That is not to say that the spatial environment is the least important driver of animal movement across all study systems, or that research should focus solely on the most important movement driver of a system. However, when studying a specific component of interest of the movement process (e.g. a spatial environmental variable), that component should still be considered properly in context of the full (*why*, *when*, *where)* movement process (Nathan et al., 2008), to avoid drawing misleading conclusions.

To showcase that misleading conclusions can follow from models in which one or more components of the movement process are ignored (‘omitted-variable bias’), we applied the same analytic pipeline to all combinations of the movement components in our case study. We identified two types of misleading conclusions that followed from models that only considered a subset of the movement process components: 1) failure to detect relationships (i.e. type II error) and 2) incorrectly detecting relationships (i.e. type I error). First, a type II error leads to overgeneralisations when drawing conclusions about the movement process. For example, wildebeest appeared to move slowest of the three species when not considering any movement process component in the model, but when movement modes were considered, the transit mode of wildebeest turned out to be faster than that of eland. While a type II error does technically not equate an erroneous conclusion (i.e., on average wildebeest did indeed move slowest), it does inhibit uncovering ecologically relevant complexities in the movement process (Spake et al., 2023). Second, a type I error leads to factually erroneous conclusions due to spurious associations between input and response variables. For example, zebra appeared to slow down substantially with increasing slope in the model with only spatial components, but with the addition of movement modes this effect disappeared completely for both modes. Apparently, mode occupancy was a confounding variable that wrongly implied a causal relationship between slope and speed (i.e. zebras were more likely to be in their encamped mode in steep areas than they were in level areas; Fig. S15). Finally, besides these two error types resulting in misleading conclusions, the omission of a movement process component that is uncorrelated to other components does not lead to erroneous conclusions or overgeneralisation, but can nonetheless still lead to simply ‘incomplete’ conclusions. For example, a model with both movement modes and temporal predictors (Fig. S12) displayed only additive effects when compared to the models that solely contained movement modes (Fig. S9) or temporal predictors (Fig. S10).

An important consideration when using our proposed analytic framework, is that we developed the daily cyclic covariate to capture both the direct, space-invariant effects of time of day on the animals’ movement properties, as well as the indirect effects of time of day through diel variation in habitat selection (given that varying environmental conditions between habitats also influence movement properties). Therefore, the daily cyclic covariate includes both a purely temporal and a spatial-temporal component. The influence of these components together on movement was large compared to that of purely spatial environmental heterogeneity (through the time-invariant tree cover and slope covariates). Future research could focus on further disentangling these temporal and spatial-temporal components further. Moreover, we see opportunities to build upon our framework with datasets that are larger in scale regarding both time and space. Namely, seasonality (e.g. through climatic variation) should come into play on top of diel rhythms as a temporal covariate when the movement data spans more months, and extra spatial environmental covariates such as vegetation quality or water availability should be added when animals roam over larger or more heterogenous areas.

To conclude, our integrative framework can be used to analyse the effect and relative importance of all components (*why*, *when* and *where* to move) of the movement process (Nathan et al., 2008). Compared to analyses that focus on a subset of these components, our approach prevents the drawing of misleading conclusions through overgeneralisations and spurious correlations. Understanding the drivers of animal movement, and ultimately of ecological phenomena that emerge from it, critically depends on considering the various components of the movement process in concert, and especially the interactions between them.

## Acknowledgments

This research was funded by the Dutch Research Council (NWO program ‘Advanced Instrumentation for Wildlife Protection’). We thank Welgevonden Game Reserve for the animal monitoring, MTN South Africa for the telecommunication and IBM for the data warehousing.

## Data availability statement

Our data and code will be made publicly available via the 4TU.ResearchData repository after acceptance of the manuscript.

## Author contributions

H.L., H.J.d.K. and J.A.J.E. conceived the ideas and designed methodology; H.J.d.K. and J.A.J.E. provided the data; H.L. and H.J.d.K. analysed the data; H.L. led the writing of the manuscript. All authors contributed critically to the drafts and gave final approval for publication.

