## Supporting Information for "Unveiling the intertwined roles of the spatial-temporal environment and behavioural modes in animal movement"

***Systematic literature search queries (Scopus, 16-03-2023)***

Spatial (400 documents):

TITLE-ABS-KEY ( spatial AND NOT temporal AND NOT ( spatiotemporal OR spatial-temporal ) AND NOT ( ( internal OR movement OR behavio* ) AND ( state OR mode ) ) ) AND TITLE-ABS-KEY ( animal AND movement AND ecology ) AND ( PUBYEAR > 2013 ) AND ( SUBJAREA ( envi ) OR SUBJAREA ( agri ) )

Temporal (72 documents):

TITLE-ABS-KEY ( temporal AND NOT spatial AND NOT ( spatiotemporal OR spatial-temporal ) AND NOT ( ( internal OR movement OR behavio* ) AND ( state OR mode ) ) ) AND TITLE-ABS-KEY ( animal AND movement AND ecology ) AND ( PUBYEAR > 2013 ) AND ( SUBJAREA ( envi ) OR SUBJAREA ( agri ) )

Modes (229 documents):

TITLE-ABS-KEY ( ( ( internal OR movement OR behavio* ) AND ( state OR mode ) ) AND NOT temporal AND NOT spatial AND NOT ( spatiotemporal OR spatial-temporal ) ) AND TITLE-ABS-KEY ( animal AND movement AND ecology ) AND ( PUBYEAR > 2013 ) AND ( SUBJAREA ( envi ) OR SUBJAREA ( agri ) )

Spatial + temporal (194 documents):

TITLE-ABS-KEY ( temporal AND spatial OR ( spatiotemporal OR spatial-temporal ) AND NOT ( ( internal OR movement OR behavio* ) AND ( state OR mode ) ) ) AND TITLE-ABS-KEY ( animal AND movement AND ecology ) AND ( PUBYEAR > 2013 ) AND ( SUBJAREA ( envi ) OR SUBJAREA ( agri ) )

Spatial + modes (95 documents):

TITLE-ABS-KEY ( ( ( internal OR movement OR behavio* ) AND ( state OR mode ) ) AND spatial AND NOT temporal AND NOT ( spatiotemporal OR spatial-temporal ) ) AND TITLE-ABS-KEY ( animal AND movement AND ecology ) AND ( PUBYEAR > 2013 ) AND ( SUBJAREA ( envi ) OR SUBJAREA ( agri ) )

Temporal + modes (24 documents):

TITLE-ABS-KEY ( ( ( internal OR movement OR behavio* ) AND ( state OR mode ) ) AND temporal AND NOT spatial AND NOT ( spatiotemporal OR spatial-temporal ) ) AND TITLE-ABS-KEY ( animal AND movement AND ecology ) AND ( PUBYEAR > 2013 ) AND ( SUBJAREA ( envi ) OR SUBJAREA ( agri ) )

Spatial + temporal + modes (47 documents):

TITLE-ABS-KEY ( ( ( internal OR movement OR behavio* ) AND ( state OR mode ) ) AND ( temporal AND spatial OR ( spatiotemporal OR spatial-temporal ) ) ) AND TITLE-ABS-KEY ( animal AND movement AND ecology ) AND ( PUBYEAR > 2013 ) AND ( SUBJAREA ( envi ) OR SUBJAREA ( agri ) )

### *Supporting figures*

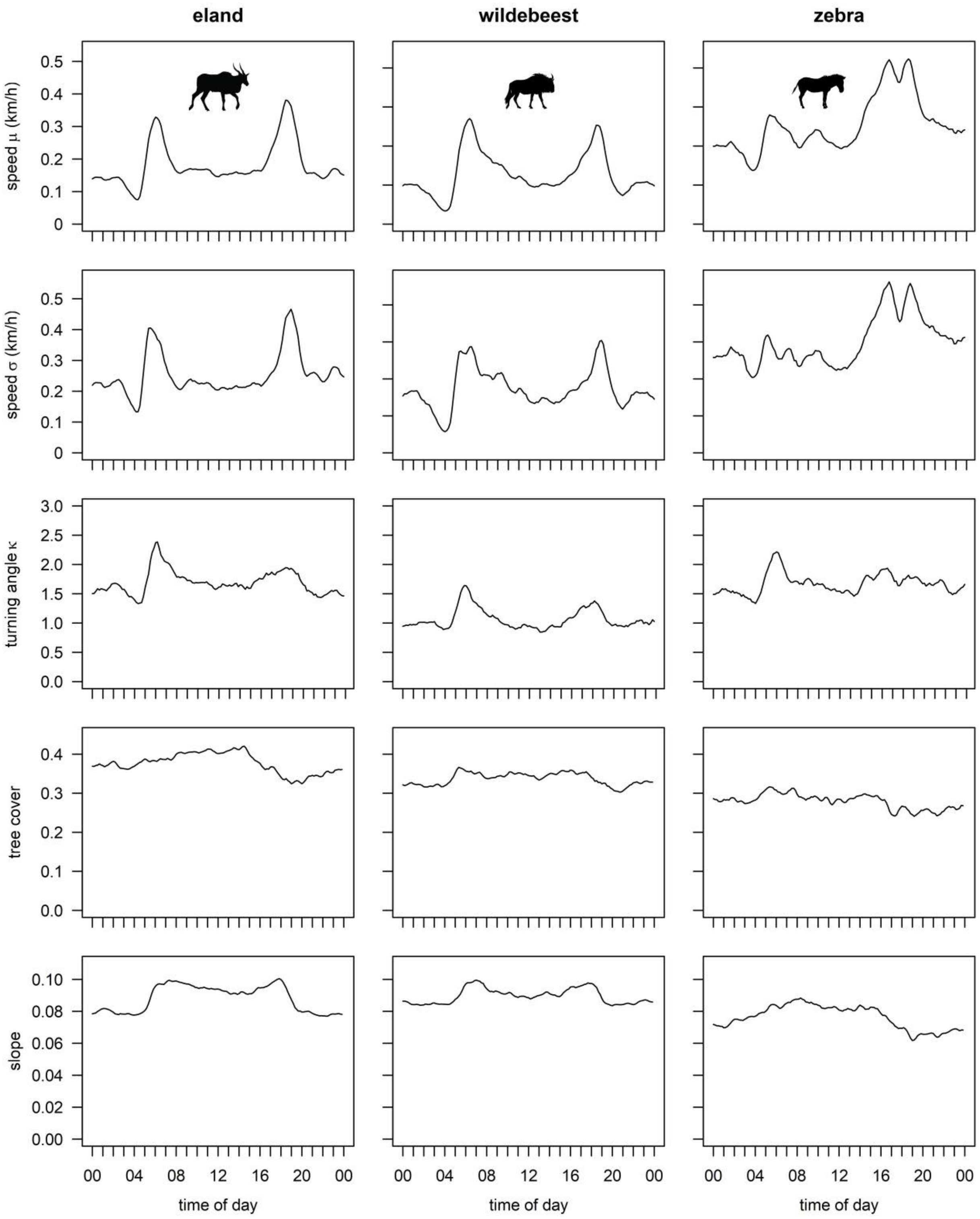


**Figure S1**. Diel patterns in speed $\mu$ and $\sigma$, and turning angle $\kappa$ (top three rows), and mean tree cover and slope (bottom two rows) for eland, wildebeest and zebra. To empirically determine the diel patterns in $\mu$, $\sigma$ and $\kappa$, we binned the GPS position data per time of day (144 bins of 10 minutes) and fitted the parameters per bin. Likewise, mean tree cover and slope were calculated within a moving window of three bins wide. For the correlations between the parameters and covariates, see Table S1.

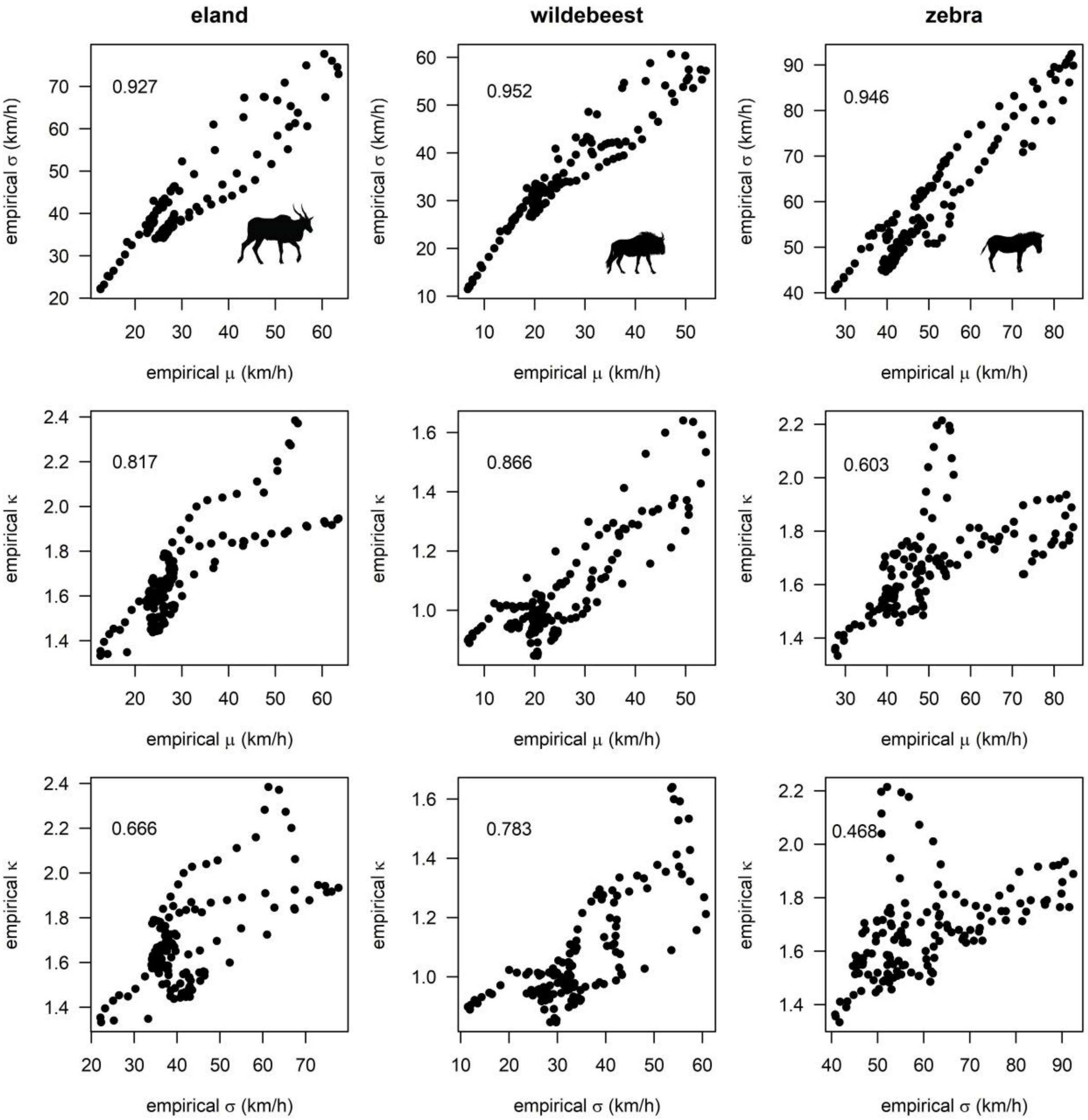


**Figure S2**. Pairwise comparisons of empirical $\mu$, $\sigma$ and $\kappa$ throughout the day for eland, wildebeest and zebra. Numbers in the graphs denote Pearson's correlation coefficients.

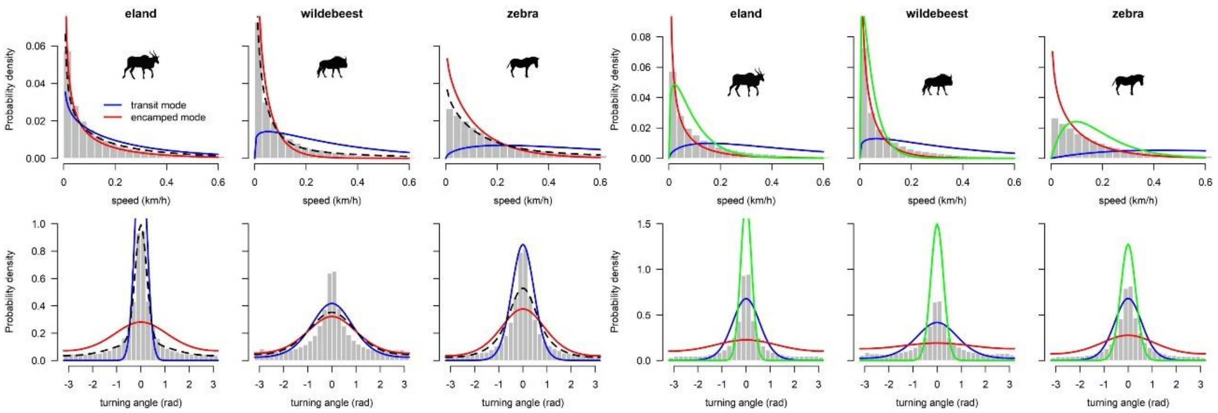


**Figure S3**. Histograms and modelled mode-dependent distributions of speed (top row) and turning angle (bottom row) for the two-mode (left) and three-mode (right) HMMs of eland, wildebeest and zebra. Different colours denote different modes; dashed lines in the left figures denote the distributions under the full model (i.e. the two modes combined). Covariates were kept constant at 0 (their mean value) for these figures.

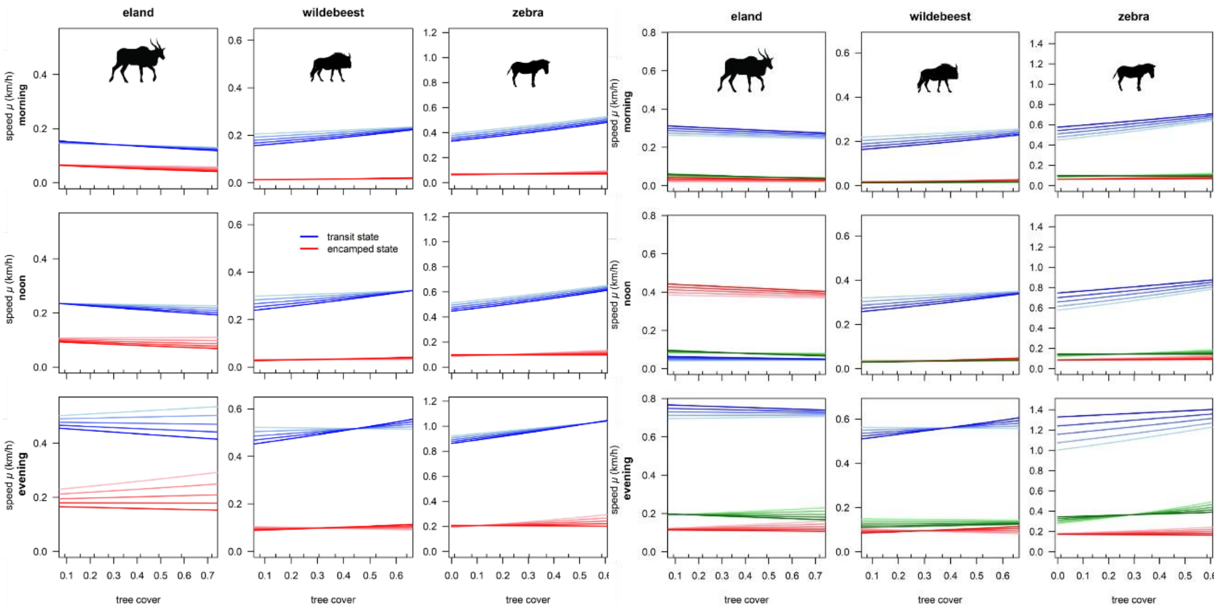


**Figure S4**. Two-mode (left) and three-mode (right) model predictions of movement speed *μ* as a result of tree cover and slope in the morning (04:00, top row), at noon (12:00, middle row) and in the evening (18:00, bottom row) for eland, wildebeest and zebra. The times were chosen to capture a dip, baseline and peak in DCC (see Fig. 3 main text). Different colours denote different modes. Colour mapping within mode colours indicates a progression of slope from 0 (light) till 0.15 (the 90th percentile of slope values in the complete dataset; dark). Small ticks at the inside of tree cover axes mark the 10$^{th}$, 20$^{th}$ … 90$^{th}$ percentile values of tree cover in the respective data sets. Note that the y-axis scales differ between species and models. Confidence intervals have been omitted for visual clarity, but they were generally small.

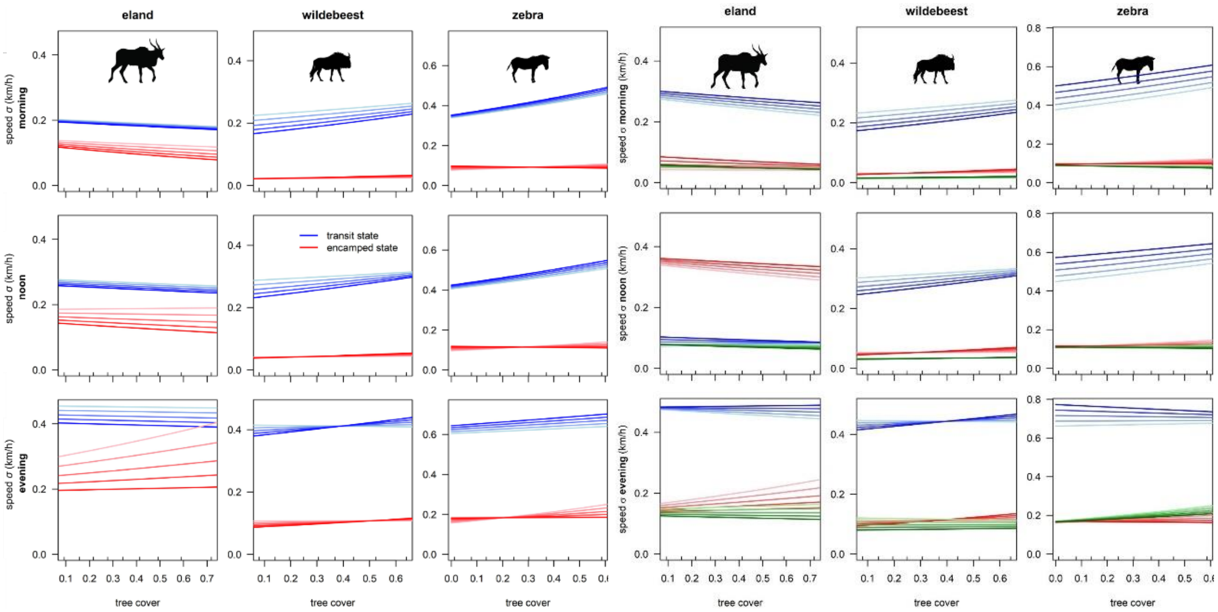


**Figure S5**. Two-mode (left) and three-mode (right) model predictions of movement speed $\sigma$ as a result of tree cover and slope in the morning (04:00, top row), at noon (12:00, middle row) and in the evening (18:00, bottom row) for eland, wildebeest and zebra. The times were chosen to capture a dip, baseline and peak in DCC (see Fig. 3 main text). Different colours denote different modes. Colour mapping within mode colours indicates a progression of slope from 0 (light) till 0.15 (the 90th percentile of slope values in the complete dataset; dark). Small ticks at the inside of tree cover axes mark the 10$^{th}$, 20$^{th}$ … 90$^{th}$ percentile values of tree cover in the respective data sets. Note that the y-axis scales differ between species and models. Confidence intervals have been omitted for visual clarity, but they were generally small.

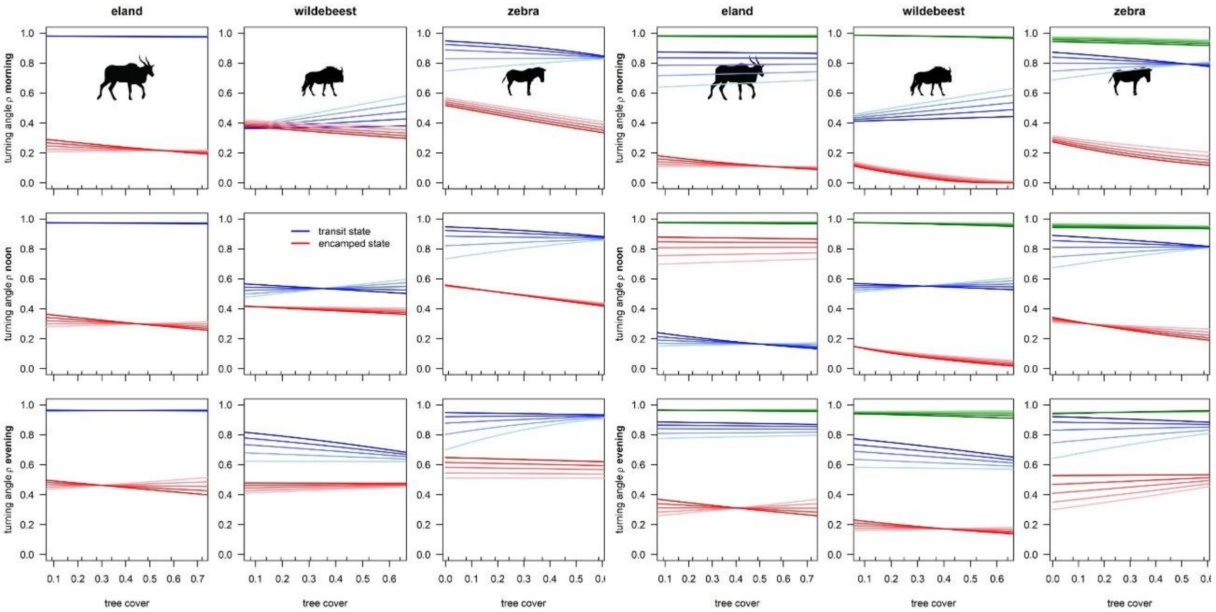


**Figure S6**. Two- (left) and three-mode (right) model predictions of turning angle concentration as a result of tree cover and slope in the morning (04:00, top row), at noon (12:00, middle row) and in the evening (18:00, bottom row) for eland, wildebeest and zebra. Concentration parameter *κ* (which ranges from 0-infinity) has been converted to *ρ* (range 0-1) in this figure for visualization purposes. The times were chosen to capture a dip, baseline and peak in DCC (see Fig. 3 main text). Different colours denote different modes. Colour mapping within mode colours indicates a progression of slope from 0 (light) till 0.15 (the 90th percentile of slope values in the complete dataset; dark). Small ticks at the inside of tree cover axes mark the 10$^{th}$, 20$^{th}$ … 90$^{th}$ percentile values of tree cover in the respective data sets. Note that the y-axis scales differ between species and models. Confidence intervals have been omitted for visual clarity, but they were generally small.

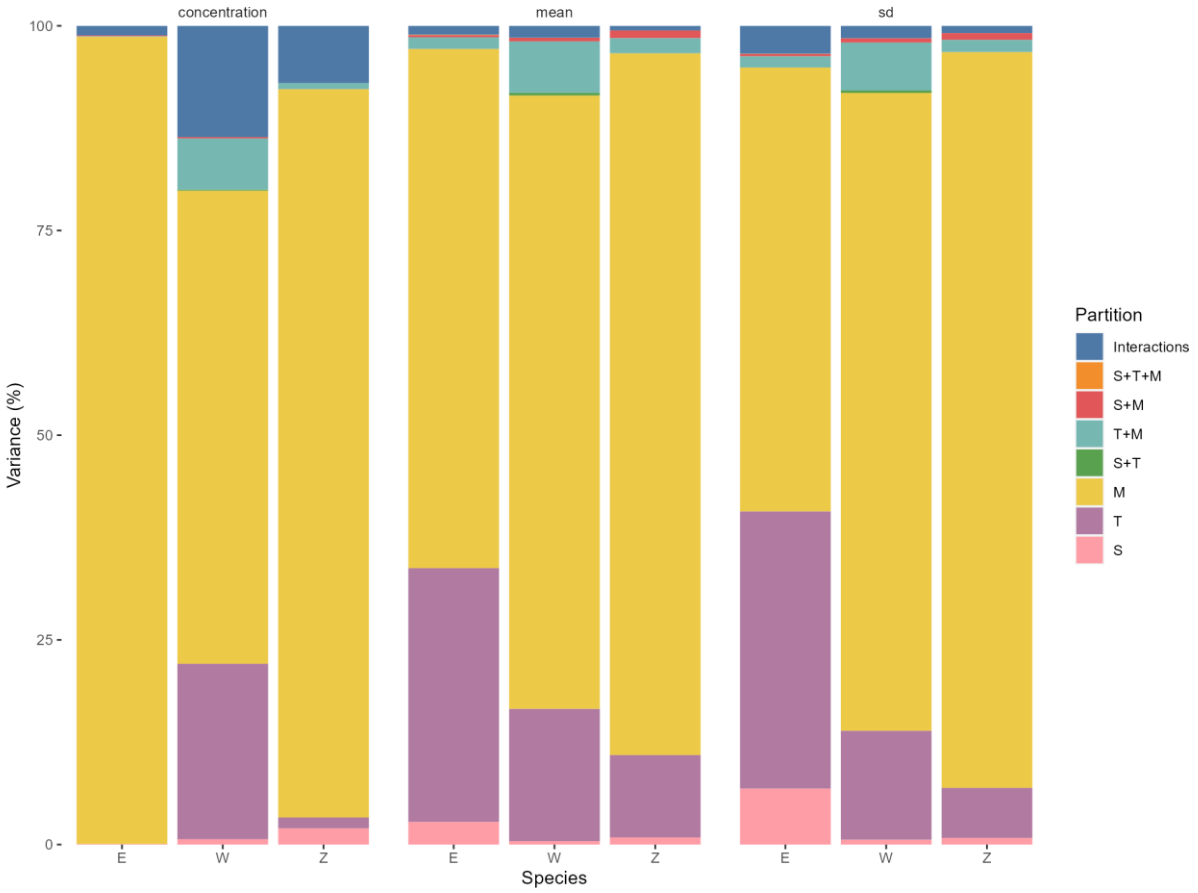


**Figure S7**. Variance partitioning, separately per movement parameter and species. E = eland, W = wildebeest, Z = zebra, M = mode, T = time, S = space.

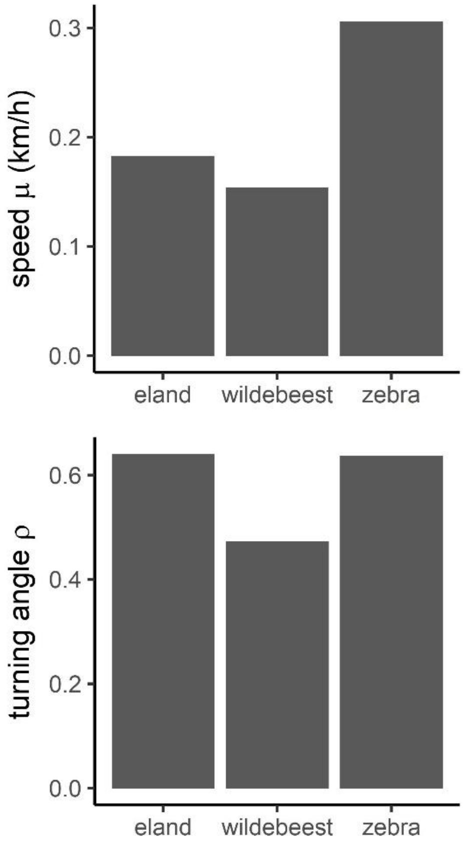


**Figure S8**. Predictions of the submodel containing neither space, time nor mode as predictors.

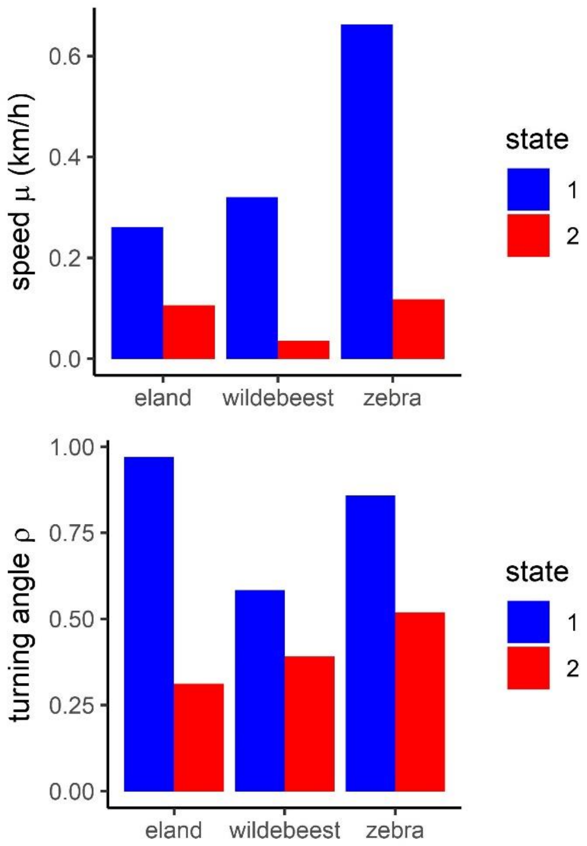


**Figure S9**. Predictions of the submodel containing only mode as a predictor.

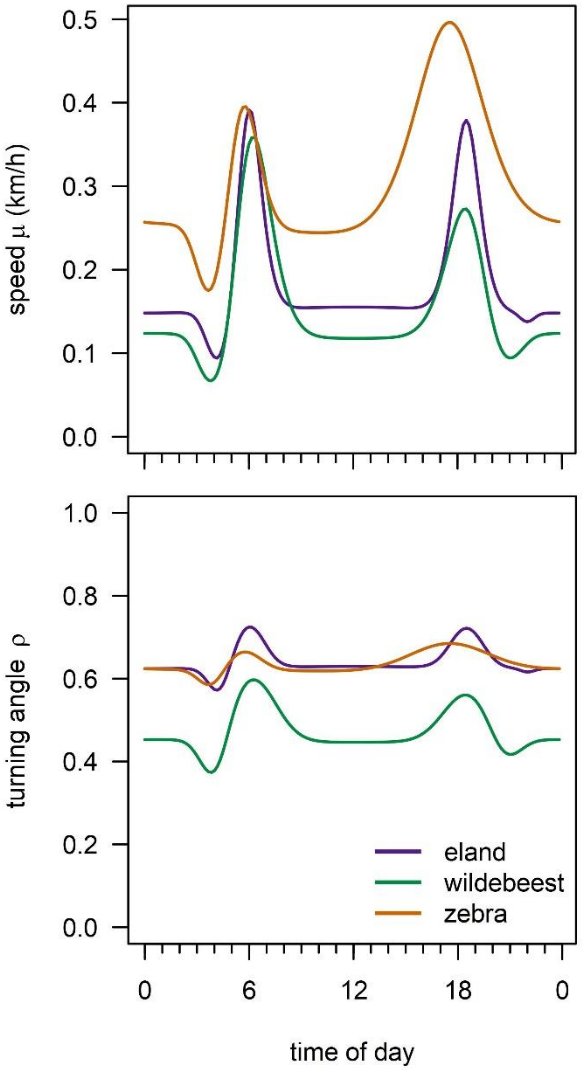


**Figure S10**. Predictions of the submodel containing only time as a predictor.

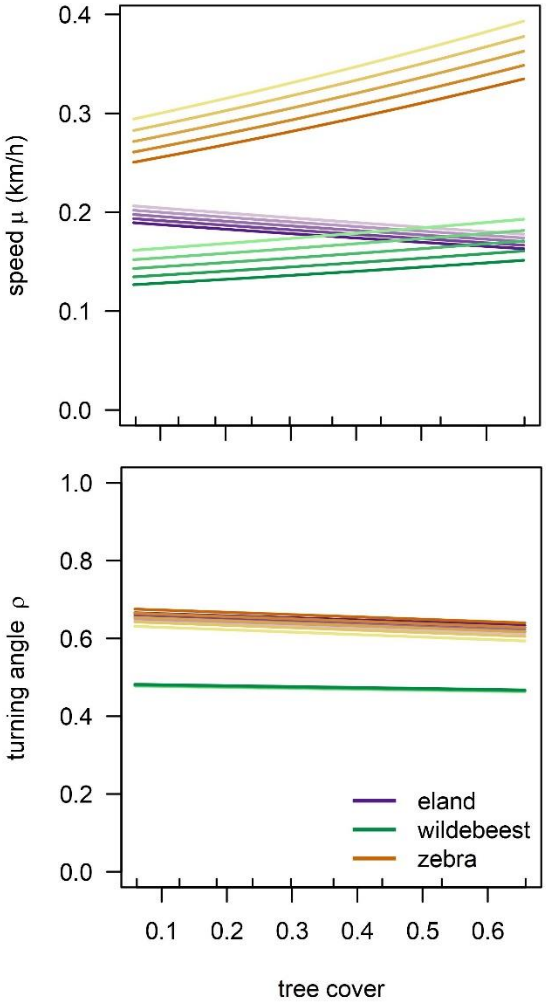


**Figure S11**. Predictions of the submodel containing only space as a predictor.

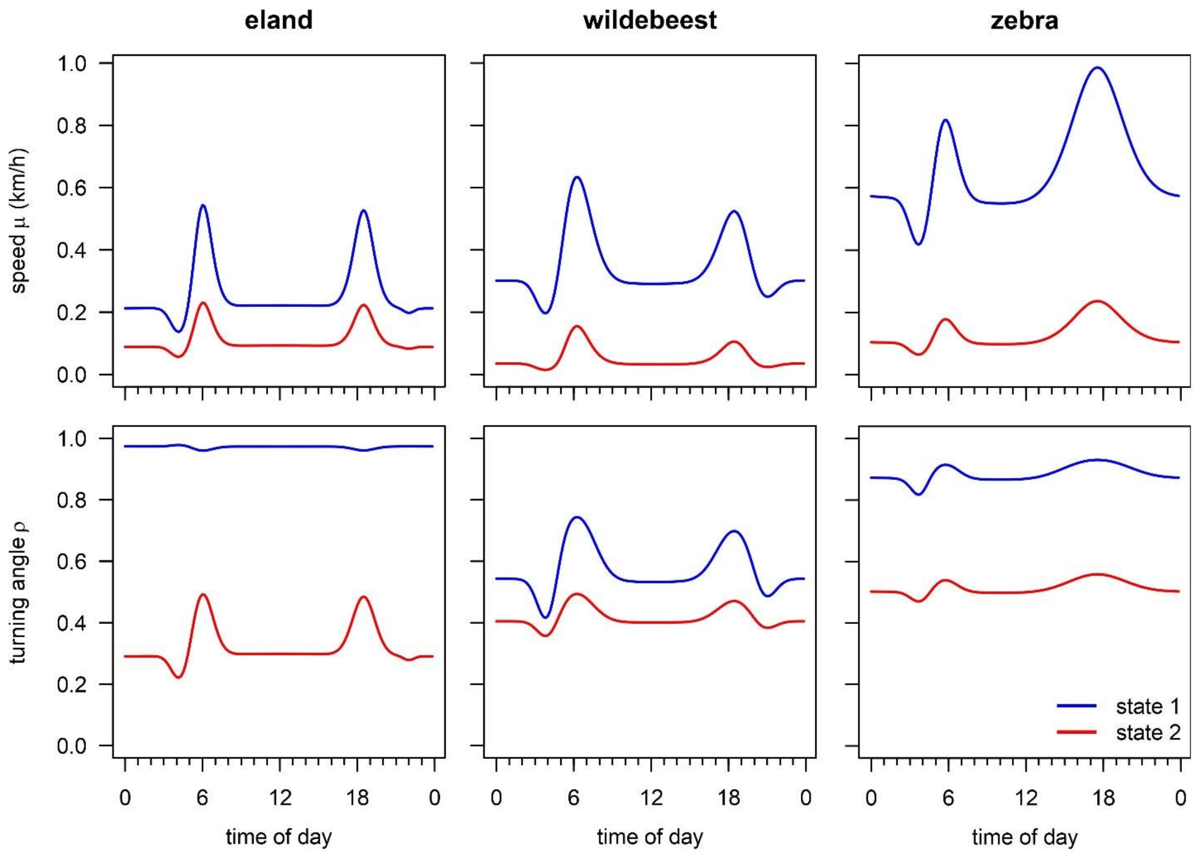


**Figure S12**. Predictions of the submodel containing only time and mode as predictors.

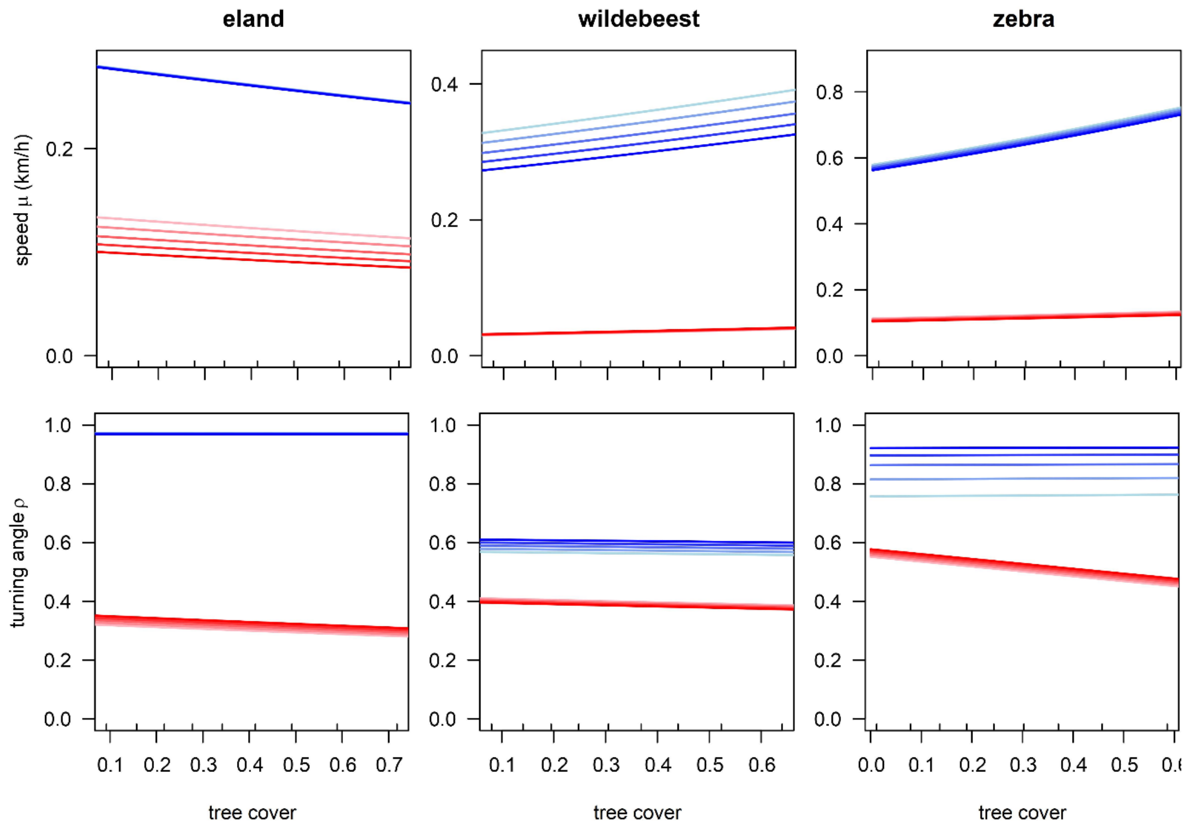


**Figure S13**. Predictions of the submodel containing only space and mode as predictors.

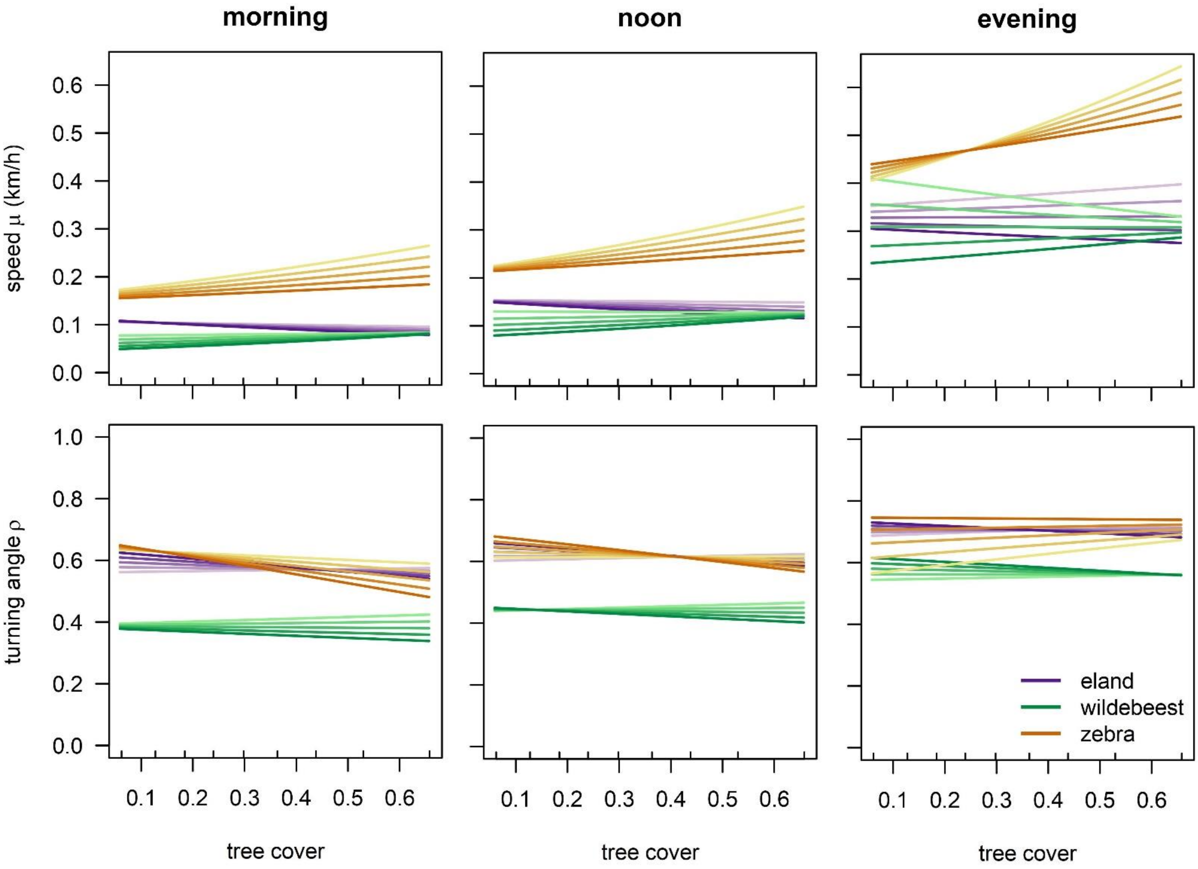


**Figure S14**. Predictions of the submodel containing only space and time as predictors.

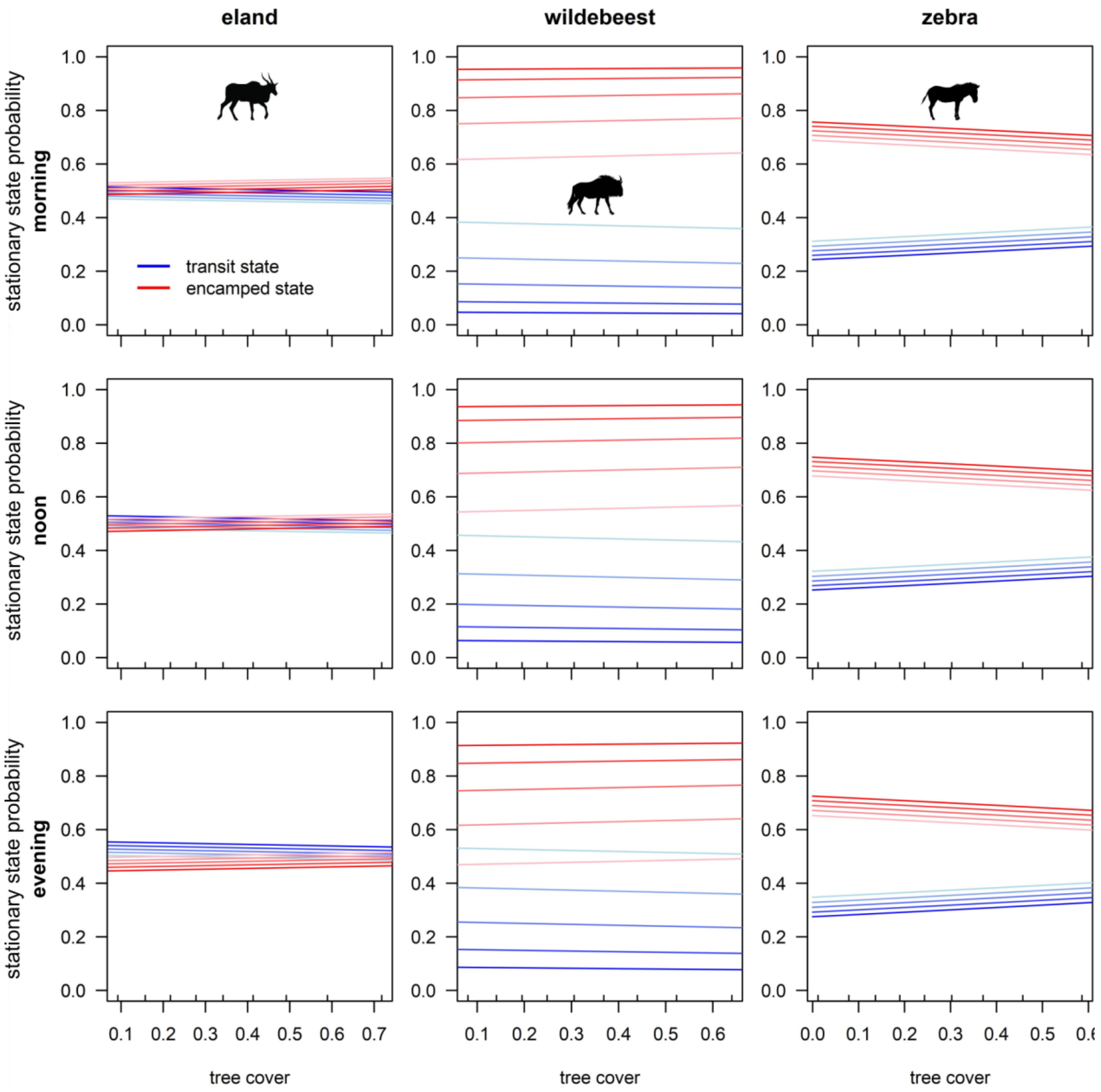


**Figure S15**. Model predictions of stationary mode probabilities as a result of tree cover and slope in the morning (04:00, top row), at noon (12:00, middle row) and in the evening (18:00, bottom row) for eland, wildebeest and zebra. Blue and red indicate the transit and encamped mode respectively. Colour mapping within blue and red indicates a progression of slope from 0 (light) to the 99$^{th}$ percentile value of slope in the respective data sets (dark); the 99$^{th}$ percentiles are 0.19, 0.17 and 0.17 for eland, wildebeest and zebra respectively. Small ticks at the inside of tree cover axes mark the 10$^{th}$, 20$^{th}$ … 90$^{th}$ percentile values of tree cover in the respective data sets. Confidence intervals have been omitted for visual clarity, but they were generally small.

### ***Supporting tables***

**Table S1**. Correlations between movement parameters *μ*, *σ* and *κ* throughout the day on the one hand and covariates tree cover and slope throughout the day on the other hand, per species. *r* = Pearson's correlation coefficient, *p* = coefficient significance.

| Species | Parameter | tree cover | | slope | |
|---|---|---|---|---|---|
| | | *r* | *p* | *r* | *p* |
| Eland | *μ* | -0.31 | <0.001 | 0.46 | <0.001 |
| | *σ* | -0.46 | <0.001 | 0.16 | 0.055 |
| | *κ* | 0.05 | 0.523 | 0.67 | <0.001 |
| Wildebeest | *μ* | 0.52 | <0.001 | 0.79 | <0.001 |
| | *σ* | 0.43 | <0.001 | 0.62 | <0.001 |
| | *κ* | 0.46 | <0.001 | 0.68 | <0.001 |
| Zebra | *μ* | -0.34 | <0.001 | -0.29 | <0.001 |
| | *σ* | -0.48 | <0.001 | -0.48 | <0.001 |
| | *κ* | 0.24 | 0.003 | 0.20 | 0.017 |
